# Generating controlled gusts using vortex rings

**DOI:** 10.1101/2021.02.09.430493

**Authors:** Dipendra Gupta, Sanjay P. Sane, Jaywant H. Arakeri

## Abstract

The control and stability of flying and swimming animals is typically determined by measuring their responses to discrete gust perturbations. For the rigorous measurement and analysis of such responses, it is necessary to generate gusts that are precise, controllable and repeatable. Here, we present a method to generate discrete gusts under laboratory conditions using a vortex ring. Unlike other methods of gust generation, the vortex ring can be well characterized and is highly controllable. We first outline the theoretical basis for the design of a gust generator, and then describe an apparatus that we developed to generate discrete gusts. As a case study, we tested the efficacy of this method on freely-flying soldier flies *Hermetia illucens*. The method described here can be used to study diverse phenomena ranging from natural flight and swimming in insects, birds, bats and fishes, to the artificial flight of drones and micro-aerial vehicles.

## INTRODUCTION

Flying animals often move in erratic environments facing discrete gusts or continuous turbulence (Etkin 1981). Such gusts are experienced when there is a sudden and sharp change in wind speed, typically encountered in the wakes of large objects or at the edge of convective disturbances (Combes & Dudley 2009; Ravi et al. 2013; Crall et al. 2017; Engels et al. 2016; Engels et al. 2019; Liao 2007). Organisms may also encounter continuous turbulence when flying in open environments or at great heights and is usually described using a statistical approach (Reynolds et al. 2016). Despite these unpredictable conditions, flying animals including insects, birds or bats are able to successfully control their flight (Combes & Dudley 2009; Ravi et al. 2013; Ortega-Jimenez et al. 2013; Victor Manuel Ortega-Jimenez et al. 2014; Ravi et al. 2016; Crall et al. 2017; Engels et al. 2016; Engels et al. 2019; Ravi et al. 2015; Ravi et al. 2020; Boerma et al. 2019). For several years, researchers from various fields have tried to quantify the locomotory abilities of diverse animals under challenging conditions. For instance, ecologists are interested in understanding how locomotory ability sets the range over which they can disperse or migrate (Chapman et al. 2011). Neurobiologists are interested in the mechanistic details of how animals sense, process and respond to perturbations to their trajectories (Dahake et al. 2018; Natesan et al. 2019). More recently, robotics engineers are interested in asking similar questions about the flight capability of their flappers and drones or swimming robots (Jafferis et al. 2019; Breuer 2019).

A key requirement for these studies is the ability to generate a well-characterized gust. To achieve this, researchers have used several methods to generate turbulence and gusts in context of natural fliers. These include grid generated turbulence (Combes & Dudley 2009; Crall et al. 2017; Ravi et al. 2015), von-Karman vortices(Ravi et al. 2013; Ortega-Jimenez et al. 2013; Ravi et al. 2016; Engels et al. 2016; Jakobi et al. 2018; Victor M Ortega-Jimenez et al. 2014; Liao et al. 2003) and compressed air jet (Vance et al. 2013; Boerma et al. 2019). Although such studies have provided keen insights into the general ways in which animals respond to turbulent gusts, the properties of the gust delivered to the animal were not precise, making it difficult to determine the state of the animals relative to the perturbation, and therefore to interpret their responses.

Here, we present a simple device that is capable of generating precise, discrete gusts using vortex rings. An impulsive flow around sharp edges when separated causes the formation of vortices that roll up into a spiral form, finally generating a vortex ring (Didden 1979; Maxworthy 1977). This vortex ring propagates with its own self-induced velocity due to vorticity concentrated in its core region. To generate precise gusts, the vortex ring is particularly effective because of its high impulse and presence of vorticity concentrated primarily in its core. We provide the necessary theoretical details to estimate flow properties of a vortex ring, followed by the description of a setup to create such vortex rings and related gusts, and the spatio-temporal characterization of their flow properties. This method can be readily adapted to a variety of contexts ranging from aerial to aquatic locomotion. Finally, as a case study, we test this device on freely-flying insects whose flight was perturbed using the gusts generated by this method.

## METHODS AND MATERIALS

### Theoretical considerations for generating vortex rings

To characterize the vortex ring as a gust source, we must first estimate its diameter and propagation speed. The most common method of generating a vortex ring is using a piston-cylinder arrangement (Fig. 1). The diameter of the rings depends on the exit diameter (D_0_) of the cylindrical tube and the distance through which the piston has moved (Auerbach 1988) (called here as stroke length) which can be equal to, smaller or larger than the exit diameter of the tube (Sullivan et al. 2008). The piston movement generates a layer of vorticity (boundary layer) in the inner wall of the nozzle to satisfy a no-slip condition. As the high-speed slug of air emerges from the nozzle, this boundary layer forms a cylindrical vortex sheet that rolls up into a spiral form, thus forming a vortex ring (Lim & Nickels 1995). In the formative stages, the vortex ring entrains the surrounding fluid as it propagates. This generates a vortex bubble (D_vb_), the diameter of which is larger than the ring (D_r_) (Fig. 1). Even after the formation process, entrainment of surrounding fluid can occur followed by subsequent detrainment, the balance between which prevents a substantial change in its diameter (Maxworthy 1972).

**Figure 1:**
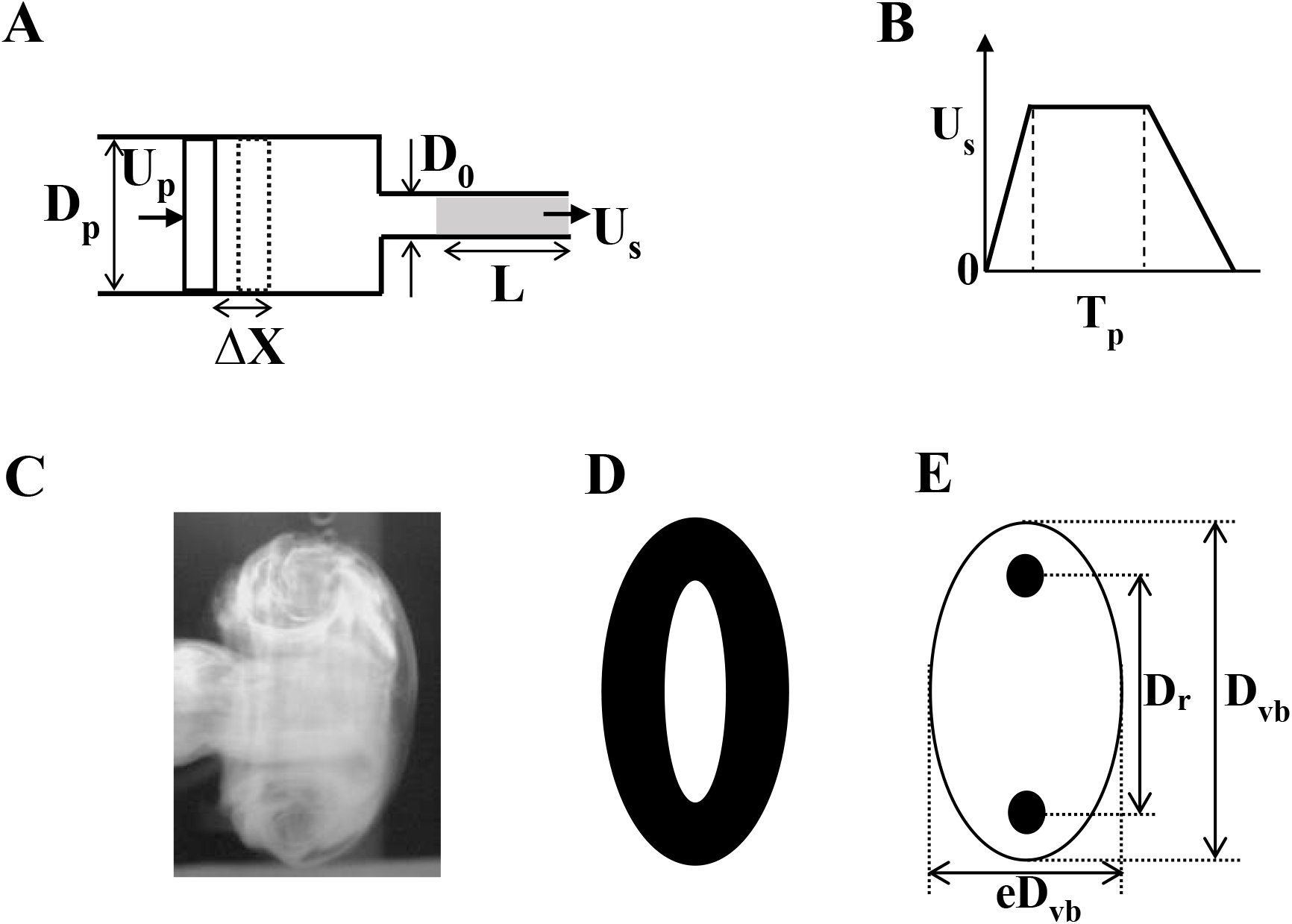
Geometric details of vortex ring and ring generator. (A) Piston-nozzle arrangement for generating vortex ring. Dashed line shows the outline of piston after it has moved by a distance *ΔX*. Grey shaded region is the length of fluid that emerges from the nozzle, called here as slug length (*L*). (B) Variation of piston velocity/slug velocity with time. *D*_*p*_ and *U*_*p*_ are diameter and velocity of piston respectively, *D*_*0*_ and *U*_*s*_ are exit diameter and velocity at the exit of the nozzle, respectively. *T*_*p*_ is the total time for which piston moves. (C) Side view, (D) isometric view and (E) assumed line diagram of side view of the ring. Black circle denotes the core of the vortex ring, in which the vorticity is concentrated. *D*_*r*_, *D*_*vb*_ and *e* denote the diameter of the ring (distance between the cores), the diameter of the vortex bubble including entrained air, and the eccentricity of the ellipsoid, respectively.

The momentum (or more precisely impulse) of the ring is determined by the momentum imparted to the surrounding fluid in the tube by the piston. It depends on the type of motion imposed on the piston, fluid viscosity, piston travel time, its radius and circulation. During vortex formation, the ring accelerates and then may rise to a constant velocity or slow down. The slowing down may be attributed to the entrainment of surrounding fluid (Reynolds 1876), entrainment followed by detrainment (Maxworthy 1972), viscous diffusion of the core (Saffman 1970) or vortex instabilities (Glezer & Coles 1990). The turbulent vortex ring, as compared to the laminar one, is characterized by a rapid growth rate of its diameter and shedding of vorticity to the wake, which causes a rapid decrease in its propagation velocity.

Thus, piston movement time (stroke time, T_p_) and slug length (L) can be designed to generate the flow properties of the ring. We provide general working relations based on simple conservation laws to achieve this objective.

#### A. Estimation of slug length (L)

Fluid is ejected out of the nozzle due to impulsive motion of the piston, the effective length of which is called slug length (Fig. 1A). Assuming the fluid to be incompressible, the volume conservation (continuity) equation gives an estimate of the slug length in relation to the velocity profile and the duration of piston motion (Fig. 1B),

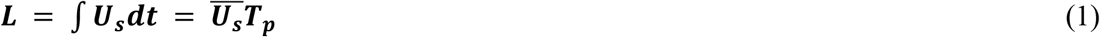

where 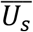 is the average velocity of the slug of fluid that leaves the nozzle exit. The slug length can thus be written as

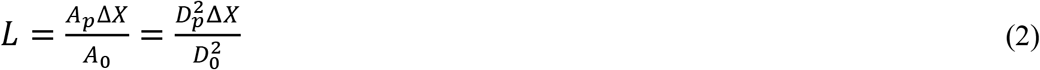

where A_p_ and A_0_ are area of piston and nozzle respectively. Equations 1 and 2 essentially say that the volume of fluid emerging from the nozzle is equal to that pushed by the piston (Das et al. 2017).

#### B. Estimation of core radius (a)

For cases where piston motion time is small, the thin vortex core assumption applies, and the core radius of the ring can be estimated (Saffman 1970) as:

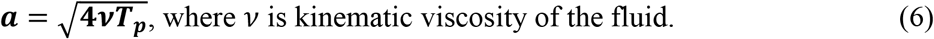

#### C. Estimation of translational velocity (U_avg_)

The velocity of the fluid ejected out of the nozzle after it is fully formed into a ring can be estimated (Saffman 1970) as

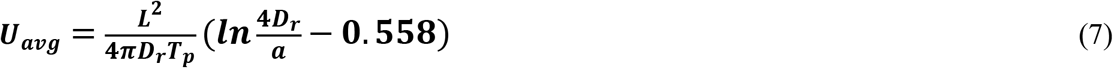

where L^2^/2T_p_ gives the estimation of circulation of the slug.

Following momentum conservation, the momentum of the slug ejected out of nozzle can be equated to the momentum of the vortex bubble, i.e.

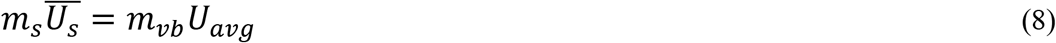

where m_s_ is the mass of fluid displaced by the piston, and m_vb_ is the mass of the ring, which includes the entrained fluid (i.e. vortex bubble). The mass of the vortex bubble is given as *m*_*vb*_ *=m*_*s*_*(1 + k)*, where k is entrained mass fraction. The value of k can be determined from the value of eccentricity, e (Sullivan et al. 2008; Loitsyanskii 1966). The entrained volume is about 20-40%total volume of the ring (Auerbach 1991; Dabiri & Gharib 2004). This implies that (8) can be written as

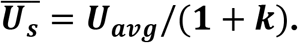

### Experimental setup

The experimental set-up consisted of a 60 cm long, 30cm square cross-section dismountable clear Perspex chamber, a 40 cm long, 3.7 cm internal diameter (D_0_) cylindrical PVC nozzle (2mm thick), a 12-inch 100W and 8 Ω speaker), a digital-to-analog converter NI-DAQ, a high voltage high current direct coupled (DC) amplifier and a high-speed camera (Miro EX4) (Fig. 2A). The large Perspex chamber served as a closed test section where we generated vortex ring. The larger dimensions of the test chamber compared to the diameter of the vortex ring reduced any effects of ambient air currents on experimental observations, and hence, aided in maintaining a still ambient fluid (Das et al. 2017). We carried the experiments in a closed room with controlled humidity at an ambient temperature of 20°C.

**Figure 2:**
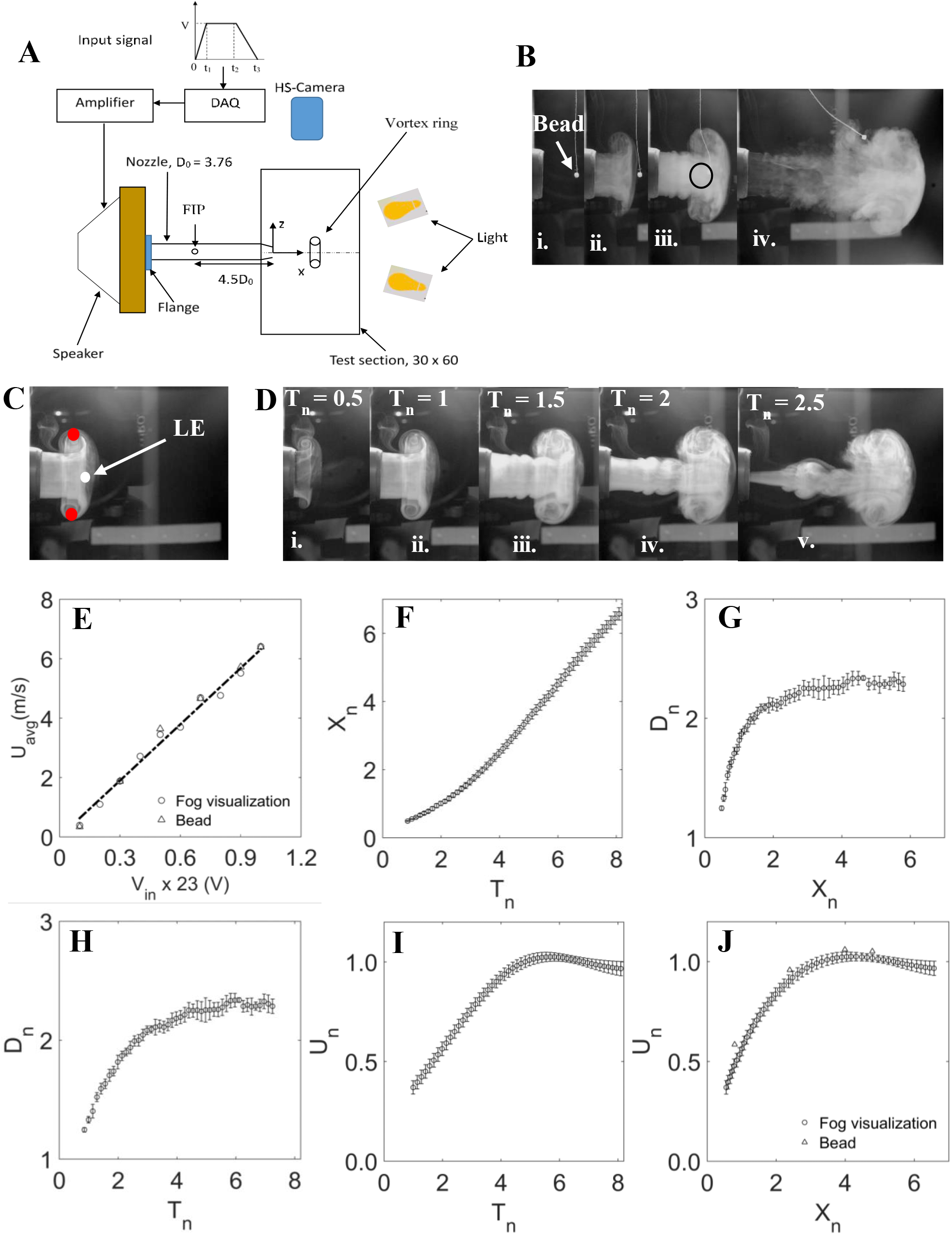
Generation and characterization of vortex ring. (A) Components of the experimental setup and their arrangement. Test chamber, Speaker, nozzle, Infrared (IR) motion sensor, Fly releasing dust (FRD), halogen lamp, cameras, Digital-to-Analog converter (DAQ) and amplifier together constitute the gust generator system. The region of interest (RoI) where the encounter of insect with the gust is desired. The input signal to the DAQ (top) is a trapezoidal wave with voltage amplitude *V*, rising time *t*_*1*_, constant duration *t*_*2*_*-t*_*1*_ and fall time *t*_*3*_*-t*_*2*_*-t*_*1*_. This signal is converted into analog form, amplified and fed to the speaker for vortex ring generation. Fog particles were injected into nozzle through fog injection port (FIP) and high-speed camera (HS-Camera) was placed horizontal to record the lateral view of the vortex ring as it propagates. All dimensions are in cm. (B-J) Flow visualization and characterization of vortex ring. (B) Effect of gust (*U*_*avg*_ = 6.4 m/s) on a freely hanging Styrofoam bead. Position of bead (i) when there is no gust, (ii) just before the gust, (iii) during gust, and (iv) after gust. The bead moves with the gust (iii) until the thread is taut. Black circle in (iii) shows the position of the bead when it is at the centre of the vortex ring. (C) White and red circles denote leading edge (LE) and extreme end of the ring. These were tracked to calculate its axial position and diameter respectively. (D) Flow visualization at different time instances for *U*_*avg*_ = 1.9 m/s. The ring propagates from left to right along X in each figure. *T*_*n*_=0 indicates the time instance when no ring is formed. (E) Average propagation velocity of the ring as a function of voltage input to the speaker. Velocity of the ring obtained using fog visualization (circles) and bead method (triangles) are in good agreement for different input voltages. Dashed line is *U*_*avg*_ = *0*.*2745V*_*in*_, R^2^ = 0.99. (F-J) Non-dimensional flow properties of vortex ring with *U*_*avg*_=6.4 m/s and vortex bubble diameter 8.6 cm measured using flow visualization and bead method. *X*_*n*_=0 indicates the centre of nozzle exit. (F) Position of ring with time. (G-H) Diameter with axial distance (G), and with time (H). (I-J) Propagation velocity showing time independency after *T*_*n*_=4 (I) and uniformity after X_n_= 3 from nozzle exit (J). Values are mean ± SD. See text for non-dimensional parameters.

Instead of a piston cylinder arrangement, we used a speaker to generate the vortex ring. The speaker was enclosed in a 40cm x 40cm x 5cm wooden chamber (driving section) on the diaphragm side, and each side of the chamber was glued with fevicol™ (Pidilite, Mumbai, India) and pin hammered to ensure that it was airtight. A 5cm diameter hole was cut at the centre of the 40cm x 40cm face of the wooden chamber to facilitate attachment of the nozzle *via* a PVC flange. A rubber gasket was used between flange and wooden chamber to eliminate any air leakage. The nozzle was sharp chamfered by an angle of 9° at the exit and smooth chamfered at entry. The speaker attached with the nozzle was then fitted to the test chamber through a 4cm hole cut on its longest side (Fig. 2A).

### Input Signal

We first synthesized a trapezoidal signal (Fig. 2A) using NI-LabVIEW, which consisted of three parts: acceleration (i.e. rise), constant velocity, and deceleration (i.e. fall). Next, we chose the time period for each portion of the signal (t_1_= 100 µs, t_2_-t_1_ = 30 ms and t_3_-t_2_-t_1_=100 ms) such that it resulted desired velocity of the vortex ring. A large deceleration time eliminates the formation of stopping vortex (Das et al. 2017) which results from abrupt stopping of piston due to separation and rolling of the secondary boundary layer induced by the primary vortex ring, on the outer surface of the tube (Didden 1979; Lim & Nickels 1995; Pullin & Perry 1980; Weigand & Gharib 1997; Shariff & Leonard 1992). We converted the signal into analog form for physical output using NI –c9263, amplified it using an in-house DC power amplifier, and fed it to the speaker, resulting in the formation of a vortex ring at the exit of the nozzle.

### Formation of vortex ring

The signal, when fed to the speaker, displaces the speaker diaphragm which imparts its momentum to the surrounding air, causing an equivalent volume of air being pushed out from the chamber into the nozzle.

Unlike conventional piston-cylinder configuration for vortex ring generation (Didden 1979; Lim & Nickels 1995; Pullin & Perry 1980; Weigand & Gharib 1997; Kumar et al. 1995; Maxworthy 1974; Auerbach 1991; Irdmusa & Garris 1987; Glezer 1988; Gharib et al. 1998; Glezer & Coles 1990; Cater et al. 2004; Sullivan et al. 2008; Maxworthy 1977; Auerbach 1987), in our experimental set-up, we have used long nozzle (11 times its diameter) so that the exit is far away from the speaker diaphragm. This eliminates the generation of piston vortex and any disturbances similar to that generated using orifice (Das et al. 2017).

### Characterization of gust

We characterized the gust using two techniques: fog visualization and Styrofoam bead method. Both the methods were carried out independently, and flow properties were characterized using each method (see *SI Video* S1).

### Fog visualization

We used fog particles to seed the flow (AntariFogger, Taiwan) for visualization of the vortex ring. The average particle size was on the order of 1-2 µm. The fog was first filled into a 500 ml wash bottle and injected through a fog injection port (FIP) on the nozzle (Fig. 2A). The port was made on the upper circumference of the nozzle, at 18 cm (4.5D_0_) away from its exit plane. The circumferential (lower, upper or sidewise) position of the port did not affect the visualization. Its longitudinal position, however, determined whether any fog particles were present at the exit of the nozzle before ring formation (i.e. background fog at nozzle exit). Keeping FIP at this distance ensures there is no any leakage of fog in the test chamber before the initiation of the diaphragm motion. A 5cm square window was hinged on the longer side opposite to nozzle. We closed the window during visualization to eliminate any effects of external air current on the ring propagation and its trajectory. After the recording, we opened the window to remove any fog inside the chamber before starting the next trial.

Figure 2B-D shows the vortex ring at different 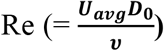 and for different time instances; *U*_*avg*_ is average propagation speed of the vortex ring (defined later), *D*_*0*_ is nozzle exit diameter and *υ* is kinematic viscosity. Re is 16000 and 4700 for the rings in Figures 2B and D, respectively. The vortex sheet emanating from the edge of the nozzle and rolling up to form a spiral and hence, the ring is visible in the pictures. The vortex ring propagated while drawing more fluid from the surrounding ambient. We did not observe any secondary and piston vortices.

### Styrofoam Bead method

In addition to fog, we used a Styrofoam bead to measure flow speeds. Because the density of Styrofoam beads (6.52 kg/ m^3^) is of an order of the density of air (see *SI Table* S1), the velocity of the bead is expected to be of nearly the same magnitude as the gust that it intercepts. This method allowed us to make point measurements of the velocity field created by the gust. We suspended the Styrofoam bead using thin sewing thread from the test chamber ceiling, such that it rested on the centreline of the nozzle exit, and intercepted the vortex ring. The bead was placed at different axial locations on the centreline of nozzle exit to measure the velocity of the bead, and hence the gust at various axial locations. One such trial for U_avg_ = 6.4 m/s is shown in Fig. 2.

Using a 12-bit CMOS camera (Phantom Miro EX4, Vision Research, Ametek, New Jersey, USA) fitted with 18-70 mm focal length lens (Nikon, Tokyo, Japan), we recorded the flow images for both these methods at 1200 fps and 50 µs exposure time. Because of the low exposure time, we additionally illuminated the background using two 1000W halogen lamps. The camera was placed to record a lateral view of vortex ring propagation in a plane perpendicular and vertical to the nozzle exit plane. The external diameter of the nozzle served as a calibration scale for the images.

We define non-dimensional time based on exit diameter of nozzle and ring average velocity, time **(t)** as 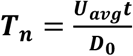. Similarly, axial distance from nozzle exit is non-dimensionalized with the exit diameter and is given by 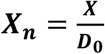. The dimensionless diameter of the ring is given by ***D***_***n***_ = ***D***_***vb/***_***D***_***0***_, where ***D***_***vb***_ is the instantaneous diameter of vortex bubble (i.e. diameter of the ring with entrained air, (see figure 1)), and dimensionless velocity of the ring is given by 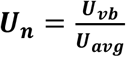 where ***U***_***vb***_ is instantaneous velocity of the vortex bubble.

### Experiments with freely-flying soldier flies

We obtained soldier flies *Hermetia illucens* from a culture housed in the National Centre for Biological Sciences. Their characteristic feature to fly towards light is one of the main reasons we chose them for study. We released a group of 5-6 flies together into a test chamber to increase the probability of them being hit by the gust. We recorded their flight motion at 4000 fps using two synchronized, high-speed cameras (Phantom VEO 640L and Phantom V611, Vision Research) for more than 80 trials, 14 of which were calibrated and digitized using MATLAB based routines-easywand5 (Theriault et al. 2014)and DLTdv7 (Hedrick 2008), respectively to measure their body and wing kinematics in response to the gust. The image data were down-sampled to 1000 fps, and then digitized and tracked head, abdomen, each wing base and tip to get their 3D position in the global reference frame. We assumed Centre of Mass (CoM) at one-third body length from the abdomen, and used it to represent the flight trajectory. X is the direction along the nozzle centreline, and Y and Z represent lateral and vertical axes, respectively.

The total velocity (velocity magnitude) of the fly is the resultant of three velocity components. A fourth-order low-pass Butterworth filter with cut-off frequency 200Hz was applied to the corresponding CoM data to minimize digitization error. We next calculated the velocity along each axis using second-order central difference scheme. We non-dimensionalized velocity by the product of body length of flies and their wingbeat frequency (*f*). Similarly, we calculated the body roll angle (*γ*) relative to the horizontal plane as the elevation angle of the vector joining the wing base and the CoM and treated it as positive if the rotation is counter-clockwise with respect to axial direction of forward flight.

## RESULTS AND DISCUSSION

The discrete gust is characterized by the spatial and temporal evolution of vortex ring and its translational velocity. We observed that the ring velocity varies linearly from 0.4 m/s to 6.4 m/s for input voltages ranging from 2.3V to 23V *(U*_*avg*_ = *0*.*2745V*_*in*_, *R*^*2*^ = *0*.*99)* (Fig. 2E), with 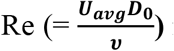 ranging from 1 x 10^3^ to 1.6 x 10^4^ respectively. Such strong dependence of the translational velocity of the ring on voltage shows that by modulating the amplitude of input signal, one can achieve vortex rings of different strengths, thus allowing control over its flow properties. The bead velocity and the average velocity of gust measured using flow visualization matched each voltage input, thus validating that bead can be used to measure such flow in air. The velocity changes in a similar fashion with time and space (distance along axis) for each voltage value. As a special case, we also discuss the ring flow properties for Re =16000 (U_avg_=6.4 m/s) (Fig. 2F-J).

The ring propagates as a quadratic function of time until T_n_ =4, after which it moves linearly until it reaches near the opposite wall of the test chamber (Fig. 2F). The ring slows down, increases its diameter, and starts getting deformed as it approaches the opposite wall. The diameter initially increases and then becomes constant with time, and can be fitted with a quadratic equation with time over the entire observation period (Fig. 2H). Because the core of the ring was not visible in all images and across all videos, we tracked the point LE to measure its axial position, and lateral extremes of vortex ring to measure its diameter (Fig. 2C). The diameter of the ring also grows as a quadratic function of space from 1.25D_0_ to 2.25D_0_ up to X_n_=3 (Fig. 2G), beyond which it attains space invariant final size of 2.3 ± 0.03 D_0_. The diameter measured here is the diameter of the vortex bubble and not the ring. Because entrained fluid generally constitutes about 20-40%of the total volume of fluid carried by tube generated vortex ring (Auerbach 1991; Dabiri & Gharib 2004) and for the present study, the eccentricity, e=0.62 and entrained mass fraction,*k*=0.57, which yields D_r_ = 0.86D_vb_

The non-dimensional ring velocity, calculated by applying second-order central difference scheme to axial position, was time-invariant after T_n_= 4 (Fig. 2I) and uniform after X_n_= 3 from the nozzle exit (Fig. 2J). The absolute value of the average speed of the ring for Re=16000 was 6.4 m/s. The representative instantaneous streamlines of the ring and its axial and radial velocity distribution at different locations and different time instances are given in *SI Figures S1* and S2. The average velocity of the ring was obtained beyond X_n_ = 3 after which it becomes nearly constant, and for the bead method, the maximum velocity of bead, which it attains in the first 5 frames of the image sequences, were considered for comparison with the average velocity obtained using fog visualization (Fig. 2E, J). We note that the maximum standard deviation in ring properties measured for three trials was less than 10%and average standard deviation less than 5%of their mean values, implying high repeatability of the measured values.

In applications, when a vortex ring is used as a gust, it should fully encompass the subject. For instance, in case of birds, bats and insects, the ring diameter should be greater than their wingspan for a head on gust. And for fish to be hit by a lateral gust, it should be greater than the body length. This allows their flight and swimming to be contained within the ring under controlled laboratory conditions. In our experiments, the ring diameter (8.5 cm) was about 4 times the fly wingspan (2.2 cm). The translational velocity of the ring should be of the order of forward velocity of the subject under consideration. While this gust velocity may hold for birds, bats and fish, in case of insects that flap their wings at higher rates, the ring velocity may need to be of the order of the wing tip velocity. In the present experiments, the ring velocity was chosen to be 6.4 m/s, which may be compared with wing-tip velocity of the fly (∼ 4.85m/s).

### Application of gust perturbation to freely flying soldier flies

To demonstrate the effectiveness of the vortex ring as a precise gust-imparting device, we subjected freely-flying soldier flies *Hermetia illucens* with head-on gusts, and measured the influence of these gusts on their flight trajectory, velocity and body angles. As mentioned above, we selected vortex ring velocity of 6.4 m/s to perturb the flies. We have chosen results from four experiments to illustrate the types of response that were obtained.

In these experiments, the flies were hit by the gust at mean axial location of X_0_ = 3.7 ± 0.38 D_0_ from the exit of the nozzle, and their lateral and vertical positions of CoM just before being hit by gust were at most 2L away from the centre of the gust (Fig. 3A, *SI Video* S2). If the flies flew to the left side of the gust, they continued to fly in the same side even after being hit by the gust. Similarly, they flew downward after the encounter with gust, indicating a possible loss in lift forces. The flies did not recover their initial vertical and lateral position after gust in any trial.

**Figure 3:**
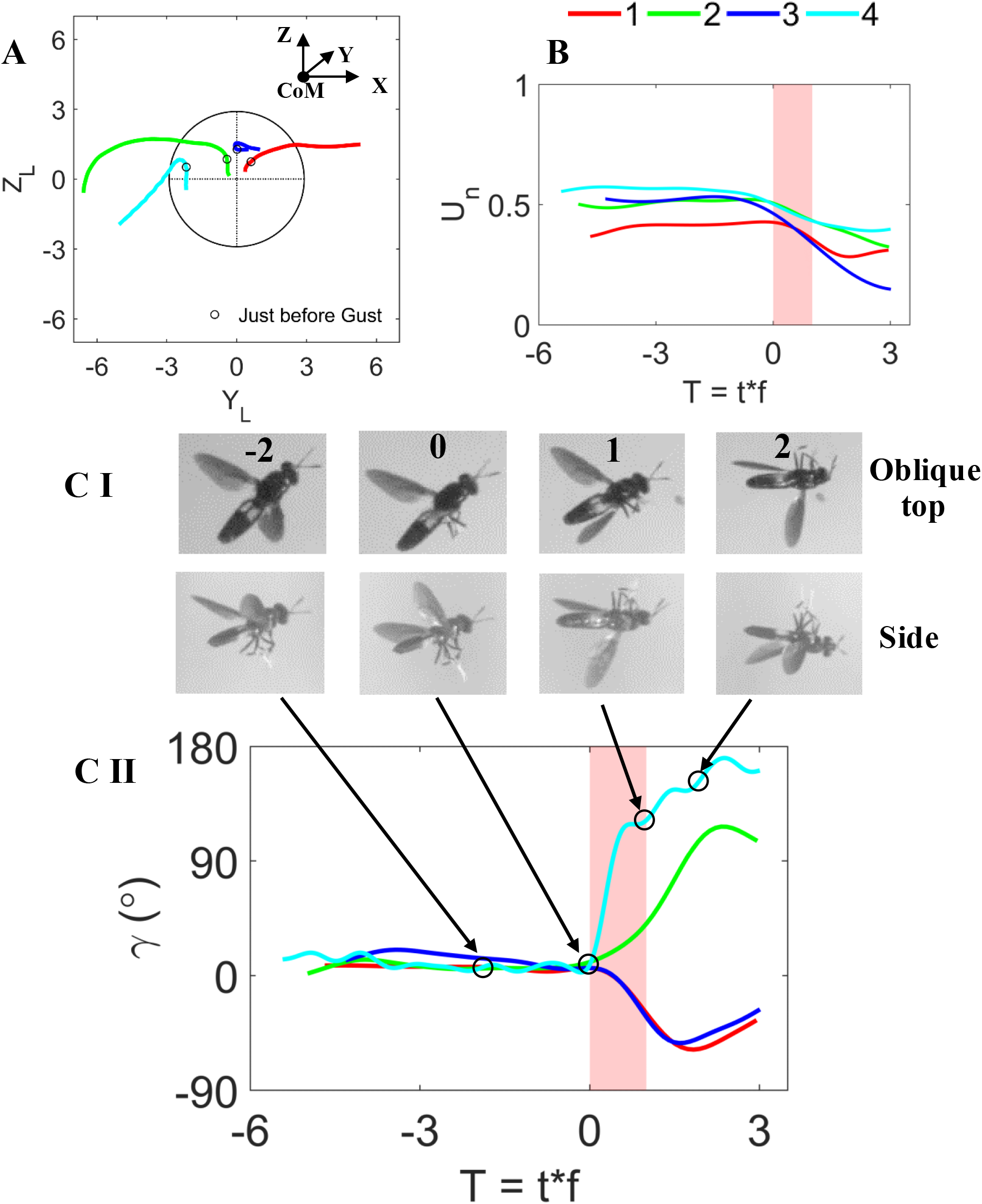
Changes in body kinematics of the black soldier flies (*Hermetia illucens)* due to the gust induced by the vortex ring in 4 different trials. (A) Trajectory in Y-Z plane normalized by the average body length (L_avg_) of flies. CoM is centre of mass. The mean axial distance where the gust hits the insects is *X*_*0*_ = 3.7 ± 0.38D_0_. Black circle indicates the front view of the vortex ring, and the intersection of vertical and horizontal dashed lines is the centre of the vortex ring. Coloured lines are the trajectories of flies for each trial represented by 1-4, and open circles on each curve denote the position of the flies just before they were hit by the ring. (B) Normalized forward speed *vs* non-dimensional time (in wing beats). Forward speed is normalized with the average speed of the flies before being hit by the ring, and time is non-dimensionalized by multiplying with the wingbeat frequency. *T*=0 indicates the time instance just before flies were just hit by the ring. Time period of gust is indicated in vertical pink strip. (C-I) Oblique top and the corresponding side views of flight sequences for trial 4 at different time instances showing distinct change in body roll angle. Number on the top row indicates the time instance of fly with respect to gust in terms of wing beats. (C-II) Change in body roll angle plotted against wingbeat. Open circles on blue curve (trial 4) denote the time instances of the fly in (C-I).

Before being hit by the gust, the forward speed of the flies was near-constant in each case. However, the gust decreased their forward speed by as much as *∼*70%maximum and *∼*30%on average (Fig. 3B). Similarly, the flies had near-zero body roll angle (<15°) before they intercepted the vortex ring, but changed by as much as *∼*160° due to gust (Fig. 3C, *SI Video* S2). The flies also responded to the ring by varying other body angles and wing stroke angle as well, the detailed results of which will be discussed in the companion paper. Thus, the observed changes in the flight trajectory, speed and body orientation show that the vortex ring can be used as a precise gust to perturb the trajectories of flies. The response and recovery of insect due to the gust will be discussed in detail in a companion paper.

## CONCLUSION

We present a method of gust generation *via* a discrete vortex ring. The ring was generated by an impulsive motion of a diaphragm of an electronic speaker. This method offers an advantage over other methods of gust generation, because the flow physics of this perturbation method is well-understood, and hence, the flow properties are highly controllable. Our method allows high repeatability and reproducibility, can be implemented at alow-cost, and has a simple mechanical design. As an example, we tested its application to study the impact of gust on insect flight, but as such this method can be used in a variety of situations and habitats. The application of the vortex ring as a gust is not only limited to insects but could potentially be extended to study birds, bats, and micro-aerial vehicles (MAVs) in air, and fish and underwater autonomous vehicles in water. We have provided relevant theoretical relations that will be useful for design of a gust generator for a specific application.

## SI FIGURE LEGENDS

**Figure S1:** Instantaneous streamlines and vorticity patch at some time after the generation of vortex ring with respect to frame moving with it, taken from Dabiri and Gharib (2004).

**Figure S2:** Velocity field of a vortex ring in a frame moving with the ring. Vortex ring dimension with core is shown in the inset. X and r are axial and radial directions respectively and O is the centre of the ring in frontal plane and R and a are the radius of the ring and its core respectively. Axial (u) and radial (v) velocity distribution along X at the centre of the ring (r = 0) (A), at r = 0.86R (B) and at r = 1.075R (C) from the centre of the ring. Radial velocity is zero at the centre and it increases away from the centre. Axial velocity decreases greatly near the core (C). In A, B and C, *Re (U*_*avg*_*R/υ)*=4.54 x 10^3^, solid lines are cubic splines fitted to the experimental data, and dashed lines denote the average translational velocity of the ring. Figures are adapted from Akhmetov(2009). Dashed line, if considered X-axis, would result in an axial velocity with respect to fixed laboratory frame, while radial velocity is invariant to such coordinate transformation. (D) Representative velocity distribution with time at any radial location, except at the centre where the radial velocity is zero.

## SI TABLE LEGEND

**Table S1:** Physical measurement of five Styrofoam beads used for flow characterization

List of symbols
|  |  |
| --- | --- |
| $a$ | *core radius of the vortex ring* |
| $A_p$ | *area of piston* |
| $A_0$ | *exit area of nozzle* |
| $D_n$ | *exit diameter of nozzle* |
| $D_p$ | *diameter of piston* |
| $D_r$ | *diameter of ring without entrainment* |
| $D_{vb}$ | *diameter of vortex bubble ( ring with entrained volume)* |
| $D_0$ | *exit diameter of nozzle* |
| $e$ | *eccentricity (= radius of minor axis/ radius of major axis)* |
| $f$ | *wing beat frequency* |
| $k$ | *entrained mass fraction (= entrained mass/mass of slug)* |
| $L$ | *slug length* |
| $m_s$ | *mass of slug coming out of nozzle* |
| $m_{vb}$ | *mass of vortex bubble* |
| $Re$ | *Reynolds number (= $U_{avg} \times D_0 / \nu$)* |
| $T$ | *no. of wing beat (= $t \times f$)* |
| $T_n$ | *non-dimensional time of vortex ring (= $t U_{avg} / D_0$)* |
| $T_p$ | *piston time* |
| $U_{avg}$ | *average propagation speed of vortex ring* |
| $U_n$ | *non-dimensional velocity* |
| $U_p$ | *piston velocity* |
| $U_s$ | *slug velocity* |
| $V_{in}$ | *input voltage* |
| $X_L$ | *axial distance non-dimensionalized by insect's body length* |
| $X_n$ | *axial distance non-dimensionalized by exit diameter of nozzle* |
| $Z_L$ | *vertical distance non-dimensionalized by insect's body length* |
| $\Delta X$ | *piston displacement* |
| $\gamma$ | *body roll angle (in °)* |
| $\nu$ | *kinematic viscosity* |
| $CoM$ | *Center of Mass* |
| $FIP$ | *Fog Injection Port* |

## Author Contribution

J.H.A., S.P.S. and D.G designed the project. D.G designed and constructed the experimental setup, carried out the experiments, processed and analyzed the data, and drafted the manuscript. J.H.A. and S.P.S. supervised the project. All authors were involved in data interpretation and preparation of final version of the manuscript.

## Acknowledgement

Funding for this study was provided by grants from the Air Force Office of Scientific Research (AFOSR) # FA2386-11-1-4057 and # FA9550-16-1-0155, and National Centre for Biological Sciences (Tata Institute of Fundamental Research) to SPS. We also acknowledge the support of the Ministry of Earth Sciences, Government of India, under project no. MESO-0034 and the Department of Atomic Energy, Government of India, under project no. 12-R&D-TFR-5.04-0800.

## References

Akhmetov, D.G., 2009. Vortex rings, Springer Science & Business Media.

Auerbach, D., 1987. Experiments on the trajectory and circulation of the starting vortex. Journal of Fluid Mechanics, 183, pp.185–198.

Auerbach, D., 1988. Some open questions on the flow of circular vortex rings. Fluid Dynamics Research, 3(1-4), p.209.

Auerbach, D., 1991. Stirring properties of vortex rings. Physics of Fluids A: Fluid Dynamics, 3(5), pp.1351–1355.

Boerma, D.B. et al., 2019. Wings as inertial appendages: how bats recover from aerial stumbles. Journal of Experimental Biology, 222(20).

Breuer, K., 2019. Flight of the RoboBee.

Cater, J.E., Soria, J. & Lim, T., 2004. The interaction of the piston vortex with a piston-generated vortex ring. Journal of Fluid Mechanics, 499, pp.327–343.

Chapman, J.W., Drake, V.A. & Reynolds, D.R., 2011. Recent insights from radar studies of insect flight. Annual review of entomology, 56, pp.337–356.

Combes, S.A. & Dudley, R., 2009. Turbulence-driven instabilities limit insect flight performance. Proceedings of the National Academy of Sciences, 106(22), pp.9105– 9108.

Crall, J. et al., 2017. Foraging in an unsteady world: bumblebee flight performance in field-realistic turbulence. Interface focus, 7(1), p.20160086.

Dabiri, J.O. & Gharib, M., 2004. Fluid entrainment by isolated vortex rings. Journal of fluid mechanics, 511, p.311.

Dahake, A. et al., 2018. The roles of vision and antennal mechanoreception in hawkmoth flight control. Elife, 7, p.e37606.

Das, D., Bansal, M. & Manghnani, A., 2017. Generation and characteristics of vortex rings free of piston vortex and stopping vortex effects. Journal of Fluid Mechanics, 811, pp.138– 167.

Didden, N., 1979. On the formation of vortex rings: rolling-up and production of circulation. Zeitschrift für angewandte Mathematik und Physik ZAMP, 30(1), pp.101–116.

Engels, T. et al., 2016. Bumblebee flight in heavy turbulence. Physical review letters, 116(2), p.028103.

Engels, T. et al., 2019. Impact of turbulence on flying insects in tethered and free flight: High-resolution numerical experiments. Physical Review Fluids, 4(1), p.013103.

Etkin, B., 1981. Turbulent wind and its effect on flight. Journal of Aircraft, 18(5), pp.327–345.

Gharib, M., Rambod, E. & Shariff, K., 1998. A universal time scale for vortex ring formation. Journal of Fluid Mechanics, 360, pp.121–140.

Glezer, A., 1988. The formation of vortex rings. The Physics of fluids, 31(12), pp.3532–3542.

Glezer, A. & Coles, D., 1990. An experimental study of a turbulent vortex ring. Journal of Fluid Mechanics, 211, pp.243–283.

Hedrick, T.L., 2008. Software techniques for two-and three-dimensional kinematic measurements of biological and biomimetic systems. Bioinspiration & biomimetics, 3(3), p.034001.

Irdmusa, J. & Garris, C., 1987. Influence of initial and boundary conditions on vortex ring development. AIAA journal, 25(3), pp.371–372.

Jafferis, N.T. et al., 2019. Untethered flight of an insect-sized flapping-wing microscale aerial vehicle. Nature, 570(7762), p.491.

Jakobi, T. et al., 2018. Bees with attitude: the effects of directed gusts on flight trajectories. Biology open, 7(10), p.bio034074.

Kumar, M., Arakeri, J. & Shankar, P., 1995. Translational velocity oscillations of piston generated vortex rings. Physics of Fluids, 7(11), pp.2751–2756.

Liao, J.C., 2007. A review of fish swimming mechanics and behaviour in altered flows. Philosophical Transactions of the Royal Society B: Biological Sciences, 362(1487), pp.1973–1993.

Liao, J.C. et al., 2003. Fish exploiting vortices decrease muscle activity. Science, 302(5650), pp.1566–1569.

Lim, T. & Nickels, T., 1995. Vortex rings. In Fluid vortices. Springer, pp. 95–153.

Loitsyanskii, L., 1966. Mechanics of Liquids and Gases, Pergamon Press.

Maxworthy, T., 1977. Some experimental studies of vortex rings. Journal of Fluid Mechanics, 81(3), pp.465–495.

Maxworthy, T., 1972. The structure and stability of vortex rings. Journal of Fluid Mechanics, 51(1), pp.15–32.

Maxworthy, T., 1974. Turbulent vortex rings. Journal of Fluid Mechanics, 64(2), pp.227–240.

Natesan, D. et al., 2019. Tuneable reflexes control antennal positioning in flying hawkmoths. Nature communications, 10(1), pp.1–15.

Ortega-Jimenez, V.M. et al., 2013. Hawkmoth flight stability in turbulent vortex streets. Journal of Experimental Biology, 216(24), pp.4567–4579.

Ortega-Jimenez, V.M. et al., 2014. Into turbulent air: size-dependent effects of von Kármán vortex streets on hummingbird flight kinematics and energetics. Proceedings of the Royal Society B: Biological Sciences, 281(1783), p.20140180.

Ortega-Jimenez, V.M., Mittal, R. & Hedrick, T.L., 2014. Hawkmoth flight performance in tornado-like whirlwind vortices. Bioinspiration & biomimetics, 9(2), p.025003.

Pullin, D. & Perry, A., 1980. Some flow visualization experiments on the starting vortex. Journal of Fluid Mechanics, 97(2), pp.239–255.

Ravi, S. et al., 2016. Bumblebees minimize control challenges by combining active and passive modes in unsteady winds. Scientific reports, 6, p.35043.

Ravi, S. et al., 2015. Hummingbird flight stability and control in freestream turbulent winds. Journal of Experimental Biology, 218(9), pp.1444–1452.

Ravi, S. et al., 2020. Modulation of flight muscle recruitment and wing rotation enables hummingbirds to mitigate aerial roll perturbations. Current Biology, 30(2), pp.187– 195.

Ravi, S. et al., 2013. Rolling with the flow: bumblebees flying in unsteady wakes. Journal of Experimental Biology, 216(22), pp.4299–4309.

Reynolds, A.M. et al., 2016. Orientation in high-flying migrant insects in relation to flows: mechanisms and strategies. Philosophical Transactions of the Royal Society B: Biological Sciences, 371(1704), p.20150392.

Saffman, P.G., 1970. The velocity of viscous vortex rings. Studies in Applied Mathematics, 49(4), pp.371–380.

Shariff, K. & Leonard, A., 1992. Vortex rings. Annual Review of Fluid Mechanics, 24(1), pp.235–279.

Sullivan, I.S. et al., 2008. Dynamics of thin vortex rings. Journal of Fluid Mechanics, 609, pp.319–347.

Theriault, D.H. et al., 2014. A protocol and calibration method for accurate multi-camera field videography. Journal of Experimental Biology, 217(11), pp.1843–1848.

Vance, J., Faruque, I. & Humbert, J., 2013. Kinematic strategies for mitigating gust perturbations in insects. Bioinspiration & biomimetics, 8(1), p.016004.

Weigand, A. & Gharib, M., 1997. On the evolution of laminar vortex rings. Experiments in Fluids, 22(6), p.447.

